# Modulation of Nur77-DNA Interactions by the Glucocorticoid Receptor

**DOI:** 10.1101/2025.10.28.685035

**Authors:** Laurens W.H.J. Heling, Kristina Kovač, Carlie J.M. de Vries, Alireza Mashaghi

## Abstract

Nuclear receptors (NRs) comprise a superfamily of (ligand-)regulated transcription factors that are pivotal in orchestrating gene networks essential for development, metabolism, and cellular homeostasis. Their activity is critical for normal physiology, and consequently, dysregulation of NR signaling is implicated in a wide array of human diseases. Within this superfamily, the orphan nuclear receptor Nur77 and the glucocorticoid receptor (GR) are key regulators that exhibit significant crosstalk, primarily antagonistic, which is crucial for modulating inflammatory and stress responses. Despite the recognized importance of their interplay, the precise molecular mechanisms by which GR modulates Nur77’s engagement with DNA remain incompletely defined. This study elucidates the direct impact of GR and its ligand, dexamethasone (Dex), on the DNA binding dynamics of Nur77. Single-molecule DNA tightrope assays revealed that Nur77 employs a three-dimensional diffusion-based search mechanism on non-specific DNA, characterized by transient interactions with two distinct dissociation kinetic profiles. GR significantly stabilizes Nur77-DNA interactions, evidenced by a shift towards longer residence times, primarily achieved by slowing the dissociation of the more transiently interacting Nur77 population. Conversely, single-molecule analysis and biochemical assays demonstrated that Dex alone markedly reduces Nur77’s overall DNA binding affinity kinetics and frequency in a sequence-dependent manner, to such an extent that accurate quantification was unfeasible. These findings delineate distinct modulatory effects of the GR protein and its ligand on Nur77-DNA interactions, providing crucial biophysical insights into their complex regulatory interplay and revealing a direct, GR-independent impact of dexamethasone on Nur77’s DNA engagement.

**Graphical Abstract:** 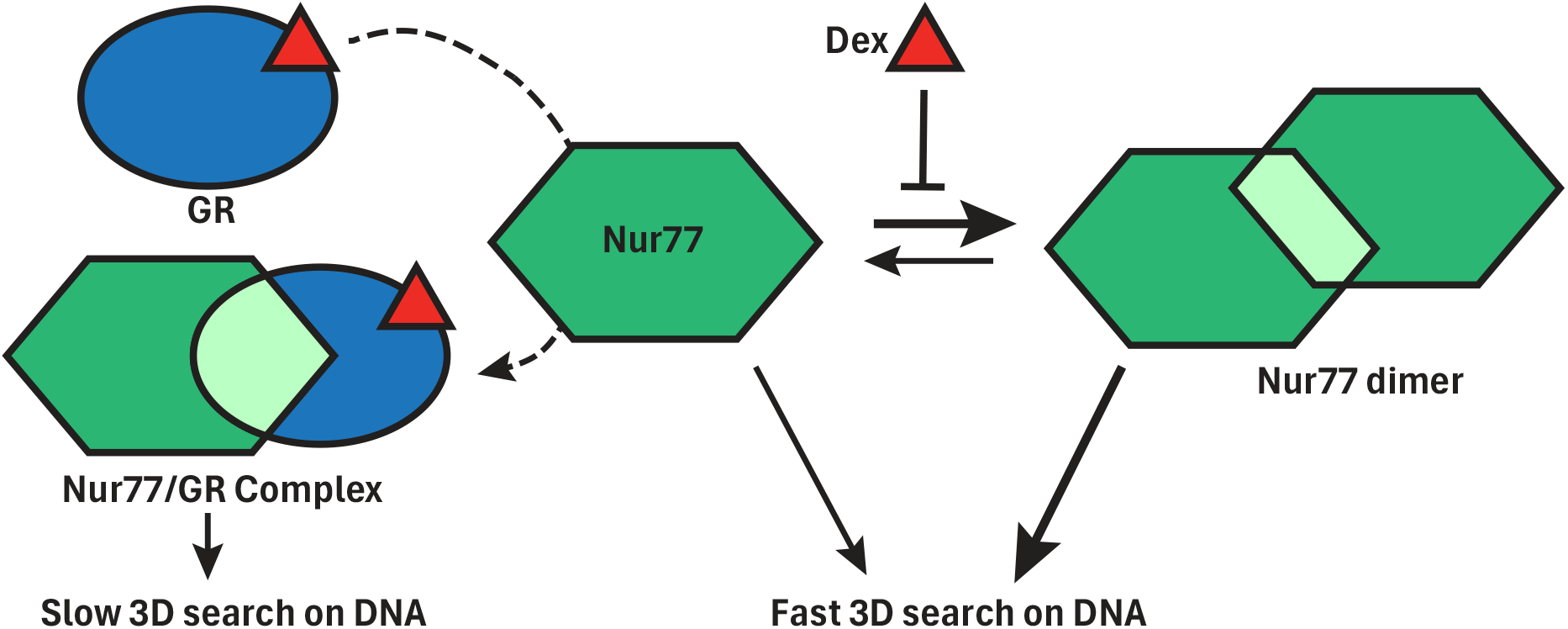

## Introduction

Nuclear receptors (NRs) constitute a superfamily of 48 transcription factors essential for regulating gene networks governing development, metabolism, cellular homeostasis, and immune response^1–3^. Dysregulation of NR signaling is implicated in numerous pathologies^4–6^. Among these, the orphan nuclear receptor Nur77 (NR4A1/TR3/NGFIB, Figure 1A-C) is a key regulator whose activity is intricately linked with that of other NRs. One of these is the glucocorticoid receptor (GR, NR3C1), which is particularly known for its activity in pathways controlling stress responses and inflammation^7,8^.

**Figure 1.**
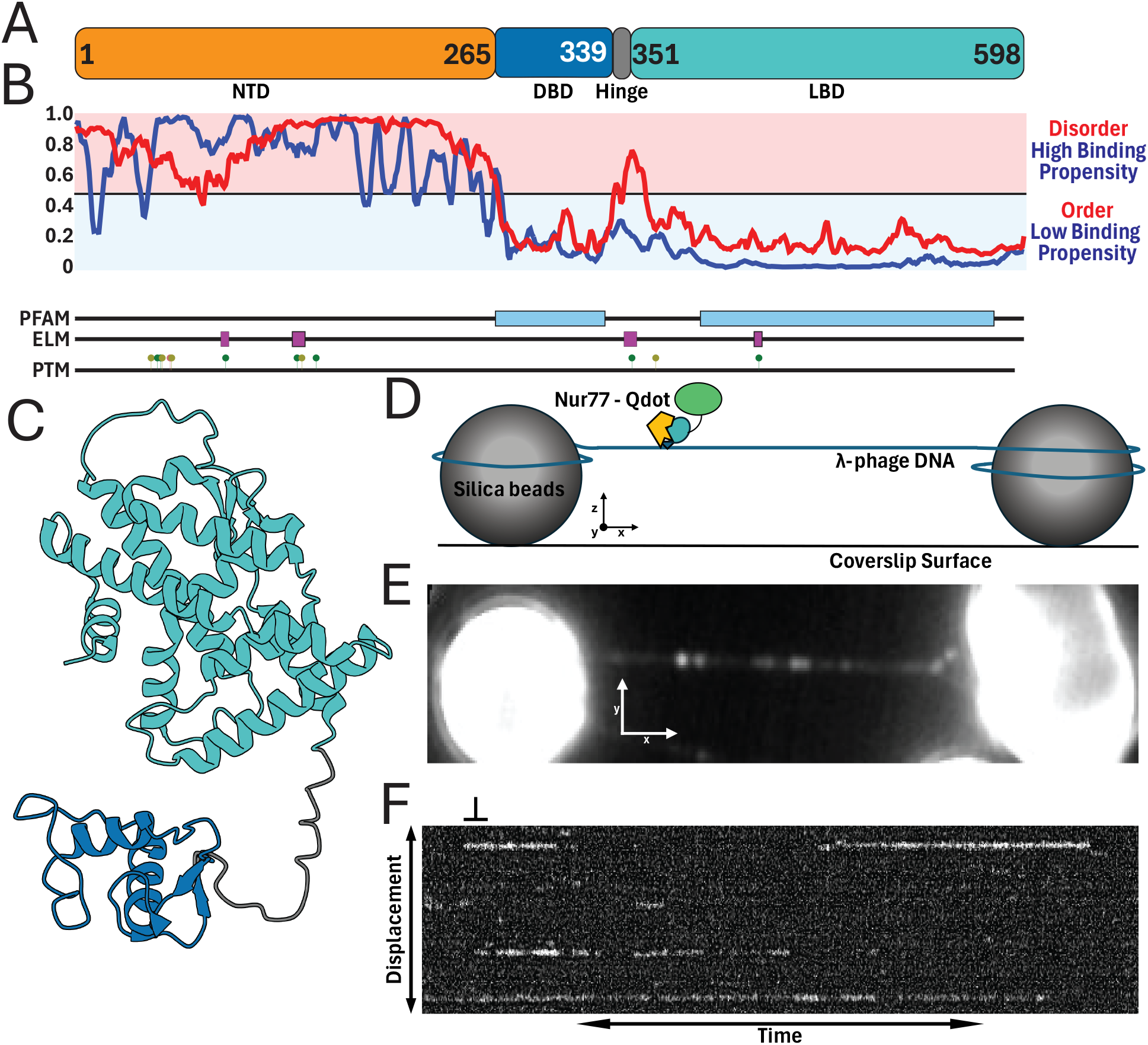
Single Molecule analysis of Nur77 DNA interactions. **(A)** Cartoon of the linear protein structure of Nur77, with the residue numbers indicating the boundaries of each region. The hinge region spans residue 340-350. **(B)** Plot generated from AIUPred^44,45^ (https://aiupred.elte.hu/). Structural analysis is shown in red, with scores above 0.5 indicating disordered regions and ordered regions have scores below 0.5. The binding propensity of the regions is shown by the blue line, with scores above 0.5 showing high binding propensity and below 0.5 low binding propensity. Underneath the PFAM domains, the recognized eukaryotic linear motifs (ELM) and low throughput translational modification sites (PTM) are highlighted. The plot is aligned with the regional delineation of the cartoon from **A. (C)** Alphafold 3 derived ribbon diagram model of human Nur77 DBD-LBD (Alphafold ID: AF-P22736-F1-v6), colour coded using Mol* viewer^46^ by canonical domain architecture as depicted in **A. (D)** Graphical representation of the DNA tightrope assay used to observe single-molecule interactions. **(E)** A maximum intensity Z-stack projection from a micrograph, illustrating a DNA tightrope stretched between silica beads with bound Nur77 molecules visible as bright spots **(F)** Kymographic transformation of a single DNA tightrope revealing Nur77 binding and releasing along the DNA over time. Scale bar represents 100 ms along the time axis and 0.727 µM along the displacement axis. Abbreviations: NTD: amino-terminal transactivation domain, DBD: DNA binding domain, LBD: ligand binding domain.

Nur77 belongs to the NR4A subfamily and exhibits unique regulatory features. While possessing a canonical NR structure (Figure 1A), the ligand-binding pocket of the ligand-binding domain (LBD) is largely occluded by bulky hydrophobic residues. Despite this, a growing list of compounds has been identified that may modulate Nur77^9–13^. However, the molecular mechanisms remain unclear and there is no consensus regarding the potential of these compounds^14^. As a consequence, Nur77 activity is thought to be governed primarily by alternative factors such as its expression level, post-translational modifications (PTMs), and crucial protein-protein interactions, rather than conventional ligand binding^15–17^. As an immediate-early gene, Nur77 expression is rapidly induced by diverse stimuli, allowing it to quickly influence cellular processes^7,16,17^. Once expressed, Nur77 engages with specific recognition sequences in the genome to modulate target gene expression. It plays versatile roles in fundamental processes including apoptosis, metabolism, and immune function^7,18–20^. Furthermore, Nur77 has been identified as a critical modulator in vascular pathobiology and has been shown to be involved in several pathologies including atherosclerosis by influencing inflammation, vascular cell fate, and lipid metabolism^21–26^. Understanding the mechanisms controlling its interactions with DNA is thus key to deciphering its diverse cellular functions.

The GR, a well-characterized steroid-activated receptor, acts as one of the key interaction partners influencing Nur77 activity^8,27^. Upon ligand binding, GR mediates widespread changes in gene expression through nuclear translocation and interactions with chromatin, coregulators, and other transcription factors, exerting both activating and potent repressive effects, notably contributing to immune suppression^28,29^. Given the overlapping physiological roles and regulatory inputs for Nur77 and GR, their direct interplay represents a significant control point.Significant crosstalk, predominantly antagonistic, exists between Nur77 and GR^17,27^. Glucocorticoids, acting via GR, can dampen Nur77 signalling both by partially inhibiting Nur77 gene induction and by directly antagonizing Nur77 protein activity^17,27,30,31^. This antagonism involves direct protein-protein interactions mediated through their respective DNA-binding domains (DBDs) ^27^. Classic examples, such as the regulation of the proopiomelanocortin (POMC) gene, illustrate this interplay where GR represses Nur77-driven transcription, potentially via recruitment to Nur77-occupied sites^17,27^. Moreover, GR activation can broadly influence Nur77’s association with chromatin, as suggested by studies on targets like the steroidogenic acute regulatory protein (StAR) gene^32^. These interactions form a critical regulatory node with consequences for hypothalamic–pituitary–adrenal (HPA) axis feedback and immune cell modulation^30,33^.

Despite the established direct interaction, precisely how this association with GR modulates Nur77’s broader engagement with DNA remains incompletely understood. Does the GR-Nur77 interaction lead to a general inhibition of Nur77’s DNA binding affinity or stability, or does it subtly reshape Nur77’s genomic interactions and transcriptional potential across diverse chromatin contexts? The mechanisms dictating how the physical interaction translates into altered DNA occupancy and function – whether through steric effects, altered protein conformation, cofactor modulation, or PTM interplay – require further elucidation^17,27,34^. A key gap lies in understanding the global impact of GR on Nur77’s DNA binding landscape. We hypothesize that the direct interaction with GR alters the fundamental DNA binding properties of Nur77, influencing its overall genomic distribution and transcriptional capacity.

This study aims to dissect the molecular consequences of the GR-Nur77 interaction on Nur77’s general DNA binding behaviour. Using a combination of single molecule DNA tightrope and biophysical approaches, we investigate how complex formation with GR modulates Nur77’s association with DNA in a simple model system. Furthermore, we mapped the impact of a potent synthetic corticosteroid, dexamethasone (Dex) on Nur77. Our results reveal direct modulation by GR and Dex on Nur77 DNA interactions, providing crucial insights on Nur77 regulation.

## Materials and methods

### Nur77 expression and purification

Sf9 cells were cultured to a density of 1.5 × 10^6^ cells/ml (500 ml) and infected with baculovirus encoding His-human Nur77. After 3 days, the cells were harvested through centrifugation. The pellet underwent 2 freeze/thaw cycles in liquid nitrogen and a water bath at 40°C. After freezing the cells for the third time, the cells were thawed in 40 mL lysis buffer (20 mM HEPES-KOH, pH 8; 1 M KCl; 10 mM Imidazole, pH 8; 100 mM L-arginine; 0,5% IGEPAL; 10% glycerol and fresh EDTA-free protease inhibitors). The viscous lysate was passed through a 21G needle 6 times to sheer the DNA (or until the lysate became less viscous). The lysate was divided into 1,5mL tubes and centrifuged for 40 min, 13k rpm, at 4°C. Ni-NTA beads (5mL) were washed twice with demineralized water and twice with wash buffer (20mM HEPES-KOH, pH 8; 1 M KCl; 20 mM Imidazole, pH 8; 100 mM L-arginine; 0,5% IGEPAL; 10% glycerol and freshly added PMSF). The lysate was applied to the beads and incubated for 3 hours at room temperature. Supernatant was removed after mild centrifugation and beads were washed 3x (4-5x column volume) with wash buffer. The beads were then transferred to a 20mL column and elution was performed with 3 ml elution buffer (20 mM HEPES-KOH, pH 8; 1 M KCl; 100 mM L-arginine; 500 mM Imidazole, pH 8; 0,5% IGEPAL; 10% glycerol; PMSF added fresh). Elution was monitored with 1x Bradford reagent (60µl 1x Bradford and 1,5µl sample because of the presence of L-arginine) (Figure S1). Proteins were aliquoted, flash frozen in liquid nitrogen and stored in -80℃.

### Microscale thermophoresis (MST)

A fluorescently labelled duplex DNA (fDNA) lacking a Nur77 recognition site was used as a substrate. The top strand was 31 base pairs long (Table S1), while the bottom strand was 32 base pairs long, creating an overhang on the 5’ end of the duplex. This overhang was filled with an aminoallyl-dUTP-ATTO-647N (NU-803-647N-S, Jena Bioscience) using DNA Polymerase I, Large (Klenow) Fragment (M0210, New England Biolabs). The reaction was cleaned up using the Monarch® Spin PCR & DNA Cleanup Kit (T1130, New England Biolabs) to remove free nucleotides. The fDNA substrate was checked for homogeneity on a 2% agarose gel. Nur77 proteins were dialysed in imaging buffer (100 mM Tris-HCl (pH 7.5), 100 mM NaCl, 20 mM MgCl_2_) overnight before being diluted from 500 nM to 2.92 nM in imaging buffer supplemented with 0.4% Tween-20 and optionally 50 nM Dexamethasone (D1756, Sigma Aldrich). These samples were mixed 1:1 with the 80 nM of the fDNA substrate in milliQ water to ensure all samples contained 40nM fDNA and Nur77 at concentrations ranging from 250 nM to 1.46 nM in 50 mM Tris-HCl (pH 7.5), 50 mM NaCl, 10 mM MgCl_2_, 0.2% Tween-20 with or without Dexamethasone., Each sample was incubated at room temperature for 10 minutes and transferred into MST capillaries (Standard Monolith Capillaries, MO-K022, NanoTemper, Germany). MST measurements were done on a Monolith NT.115 instrument (NanoTemper, Germany) at 40% LED power and medium MST power. Total measurement time was 30s, with 5s laser off, 20s laser on and 5s laser off. F_norm_ values were evaluated after 10s of laser on and normalised against baseline (DNA alone).

### Single-molecule DNA tightrope assay

The DNA tightrope assay was described previously in detail^35,36^. In short, a custom microfluidic flow chamber was made by combining a standard microscope slide with two holes drilled in it, a double-sided tape gasket and a silanized coverslip. Two tubes connected to a syringe on a peristaltic pump (World Precision Instruments, AL1000-220) and a custom Eppendorf tube enabled controlled movement of reagents through the flow chamber. To reduce non-specific surface interactions of proteins, DNA or fluorophores, the flow chamber was blocked by overnight incubation in mPEG buffer (25 mg/ml mPEG5000 in 250 mM NaHCO3, pH 8.2) followed by incubation overnight in ABT buffer (10 mg/ml BSA, 0.1% Tween-20 & 0.1% NaN_3_). After careful washing of the flow chamber with buffer, 5 μm poly-L-lysine silica microspheres (SS05003, Bang’s Laboratories) coated with poly-L-lysine (p5899, Sigma Aldrich) are deposited on the surface and λ-phage DNA (500 ng) is flowed back and forth through the chamber at a constant velocity (300 μL/min) for 40 minutes, elongate the DNA molecules and forming single molecule DNA tightropes between the beads.

Proteins were labelled with fluorescent Quantum dots (Qdot). The his-tag located on the Nur77 was tagged through a Qdot antibody sandwich. Nur77 (1 μM) was dialysed against the imaging buffer (described in the MST section) overnight. An equimolar concentration of Hexa-His Primary Antibody (10001-0-AP, Proteintech) was added to the protein and incubated on ice for 30 minutes. A F(ab’)2-Goat anti-Mouse IgG (H+L) Secondary Antibody with a 525 Qdot (Q-11041MP, Invitrogen) at was then added to the protein:primary antibody solution at a 3:1 excess (to ensure single labelled Qdots) and incubated for 30 minutes on ice. Labelled proteins were diluted in imaging buffer, and prepared for introduction into the tightrope assay. The final concentration of Nur77 was 1 nM, which ensured single molecule resolution with a limited number of molecules that can interact with each DNA tightrope. The imaging buffer was supplemented with 10 mM DTT, which we found reduced “blinking” of the Qdot^37^, a phenomenon where the emission of the Qdot is temporarily reduced. We incubated Qdots, without protein, into the DNA tightrope assay to exclude direct DNA binding by the Qdots (data not shown).

For experiments with GR, commercially available recombinant full-length human GR (ab82089, Abcam) was activated with dexamethasone to avoid aggregation. This sample was filtered to reduce any present aggregates and aliquoted. 1 nM of dexamethasone activated GR was introduced into the flow chamber via the reservoir tube. For experiments with dexamethasone, 25 nM dexamethasone was introduced to the flow chamber through the reservoir.

### Super resolution fluorescence microscopy

Single-molecule imaging was performed on a home-built wide-field setup, based on an Axiovert S100 (Zeiss, Germany) inverted microscope equipped with a 100x 1.4NA oil-immersion objective (Zeiss, Germany) at room temperature (20 °C). The sample was excited by a 488nm laser (Sapphire CDRH, Coherent Inc., USA). The intensity and timing were set through an acousto-optic modulator (AOTFnC-VIS, AA-OptoElectronic, France) such that the intensity was set at 1 kW/cm^2 and the illumination time at 10 ms per frame. The light was detected through a dichroic/emission combination (Di01-R405_488_561_635/zet405_488_561_640m, Semrock, USA). The signal of individual dye molecules was captured on a sCMOS camera (Orca Flash 4.0V2, Hamamatsu, Japan). The comparison of the single molecule fluorescent signal to background yielded a signal-to-noise ratio of 14 and were spatially distributed according to the microscope’s point-spread function (440 nm FWHM), allowing for localization of individual fluorophores with sub-30 nm precision^38^.

### Data analysis

For single-molecule microscopy, three independent flow cells were prepared per condition. When a suitable tightrope was detected, 60 second videos were recorded. The videos were converted into kymographs using ImageJ^39^. To identify tightrope positions between beads, videos were projected as Z-stacks using maximum intensity projection. A 25-pixel rolling ball radius background subtraction was applied to the kymographs, to average the intensity of bright and dim binding events. Kymograph projections allow the identification of different DNA search mechanisms: protein sliding along the DNA to search for its target sequence will appear as movement along the y-axis over time, while a 3-dimensional search mechanism is represented by horizontal streaks. For the lifetime analysis, molecules exhibiting binding exceeding the video length were excluded. To mitigate bin size bias, attached lifetimes were plotted as cumulative frequency histograms and fitted. The necessity of single versus double exponential fits was assessed via F-test. Data fitting was performed with a custom-built Python code, using numpy^40^, pandas, matplotlib^41^, seaborn^42^ and scipy^43^ libraries. Additional error analysis was performed in Origin2024 to determine fitting errors using the parameters from Python fits.MST data was normalised against DNA alone and plotted using Microsoft Excel. The fitting and *k*_*D*_ calculations were performed using a custom Python script.

## Results

### Nur77 performs a fast-paced 3D search on DNA

Transcription factors like Nur77 may use a number of search mechanisms to find their target DNA sequence *in vivo*^*47*^. To directly visualise the search mechanism of Nur77 for a sequence in real time, we employed the *in vitro* DNA tightrope assay. In this assay, individual DNA molecules are suspended between 5 µm silica beads deposited on a PEGylated surface of a custom-built microfluidic chamber (Figure 1D)^35,36^. This allows for the direct observation of protein-DNA interactions where the protein has full accessibility to the DNA while improving signal-to-noise ratios. We labelled individual Nur77 proteins with a bright photostable Qdot via an anti-His-tag antibody conjugation. We incubated Nur77 with lambda-phage (λ) DNA, a long DNA substrate ideal for the tightrope assay (Figure 1E). Two different recognition sequences have been identified for Nur77 *in vivo*, the NurRE (TGATATTTn_6_AAATGCCA), to which Nur77 binds as a homo or heterodimer (with other NR4A members), and the NBRE sequence (AAGGTCA) which is bound by Nur77 as a monomer. λ DNA lacks the full canonical binding NurRE but does contain a few half NurRE and NBRE sites. We observed interactions within a 60 second time window. Videos of Nur77 interactions with DNA were transformed into kymographs (Figure 1F), from which different types of diffusion behaviour can be characterised ^36^.

Our data revealed that Nur77 associated with DNA and did not exhibit any sliding behaviour along the DNA within the spatiotemporal limits of our setup (<30 nm, ∼88 bases, 10 ms). The wide range of observed binding lifetimes (n = 809), ranging from 0.03 to exceeding 15 seconds, aligns with the stochastic nature of molecular dissociation events, and is likely indicative of a kinetically heterogeneous population. Together these findings suggest that Nur77 probes DNA for its target sequence using 3-dimensional diffusion dynamics.

### GR shifts lifetime equilibrium to longer lived states

Upon introduction of equimolar concentrations of Dex-activated GR, the data indicate a discernible shift towards longer lifetimes. The average lifetime observed for Nur77 with (Dex activated) GR present was 1.32 ± 0.06 seconds, which is markedly higher than the average of 0.98 ± 0.06 seconds for Nur77 alone. The trend towards longer durations is further supported by an examination of the full range of observed lifetimes (Fig. 2A). This distributional shift is further characterized by a notable reduction in the number of events where Nur77 resides on the DNA less than 0.3 seconds, and a concomitant enrichment of species persisting in the 0.5 to 2-second range. Consistent with the right-skewed distributions typical of single-molecule decay, the presence of (Dex activated) GR extended the tail of longer-lived molecules.

**Figure 2.**
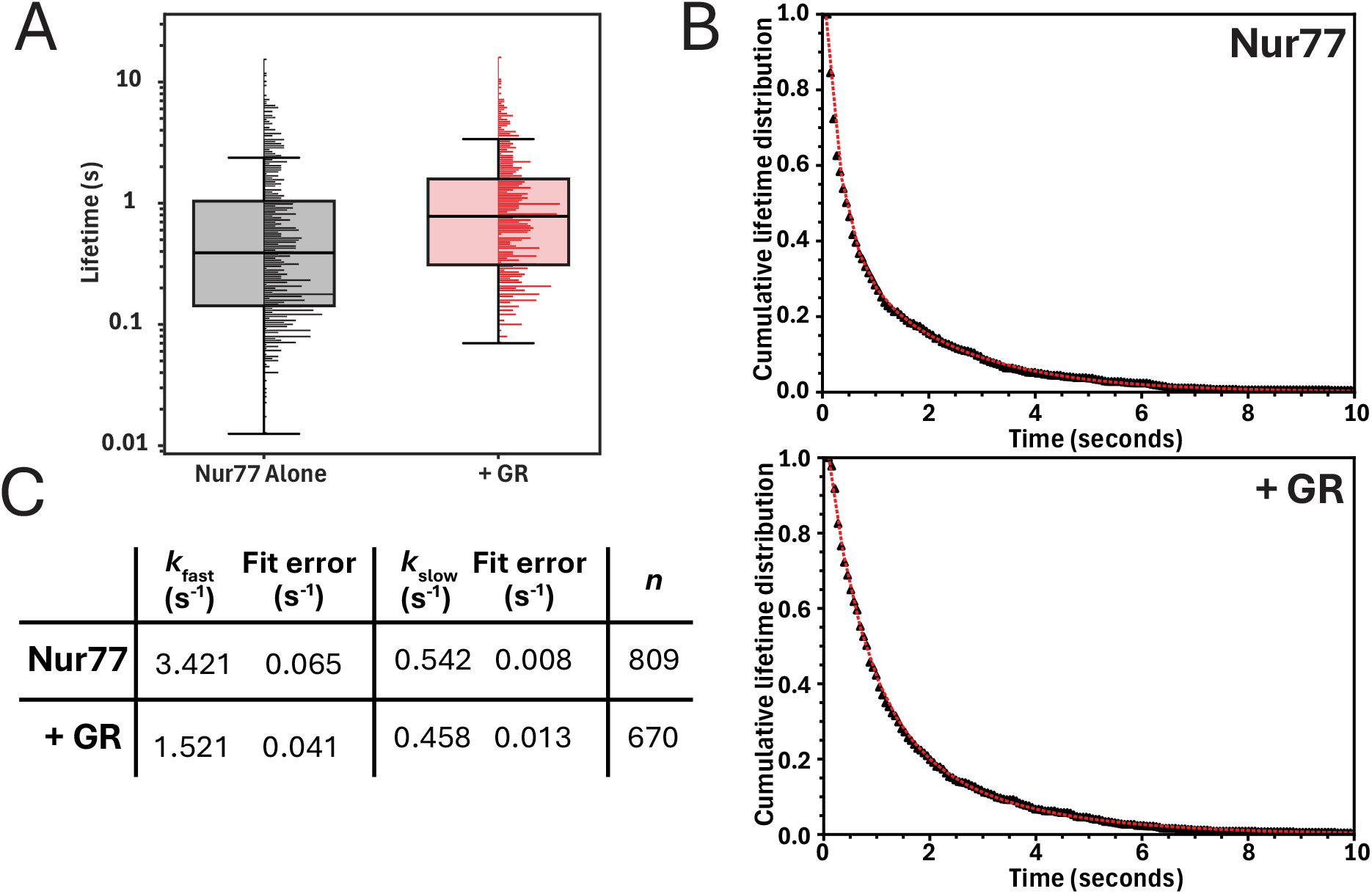
Nur77/DNA interactions dynamics are modulated by GR. **(A)** Boxplot with accompanying normalized histograms of the molecular lifetimes (in seconds) reveal that (Dex activated) GR significantly increases Nur77 lifetimes on DNA (Mann-Whitney U test *p* < 0.001). **(B)** Top: Cumulative lifetime distribution of Nur77 interacting with DNA alone. Bottom: cumulative lifetime distribution of Nur77 in addition of Dex-activated GR interacting with DNA. Experimental data is represented by black triangles with double exponential fits (red dotted line), **(C)** Table displaying the dissociation rate constants (*k*_fast_ and *k*_slow_) extracted from the double exponent. The fit error was determined using Origin2024 using the fitting parameters. *n* represents the number of observations from three separate flow chambers.

Collectively, these data demonstrate that activated GR alters Nur77 molecular dynamics on DNA. This is observed not merely as an extension of a few long-lived outliers but rather a comprehensive shift in the entire lifetime distribution.

The dissociation of proteins on DNA can be modelled as a Poisson process^48^. To further elucidate the heterogeneity in residence lifetimes induced by GR, and to extract the rate constants, we plotted the data as cumulative lifetime distributions (Figure 2B). Fitting the data yields the dissociation rate constant (k_off_). If a system has multiple intermediates, the number of terms fit to the distribution can indicate the number of intermediates. Our data fit best to a double exponential function suggesting the presence of two kinetic intermediates for both the Nur77 alone (Figures 2B-C and S2) as Nur77 with GR conditions (Figure 2B-C and S3). For Nur77 alone, approximately two-thirds of events (63.3%) exhibited short dwell times on DNA, with a *k*_off_ of 3.421 ± 0.065 s-1, while the population of longer-lived interactions (36.7%) resided at the DNA had a *k*_off_ of 0.542 ± 0.008 s-1. (Figure 2C). In the presence of GR, the population distribution between short- and longer-lived populations were similar to Nur77 alone (63.5% and 36.5% respectively), yet the rate constants decreased. While the longer-lived populations demonstrated only a moderate decrease with a *k*_off_ of 0.458 ± 0.013 s^-1^, the short-lived population the rate constant was less than halve with 1.521 ± 0.041 s^-1^.

### Dexamethasone disrupts Nur77 DNA binding

Given that Dex, a potent synthetic glucocorticoid, was added to ensure GR was in its active form, we conducted additional experiments to understand the direct impact of Dex on Nur77 binding kinetics. First, we compared DNA binding of Nur77 with and without Dex using microscale thermophoresis (MST). This technique quantifies molecular interactions by detecting changes in the diffusion of fluorescent molecules within a temperature gradient (Figure 3A).

**Figure 3.**
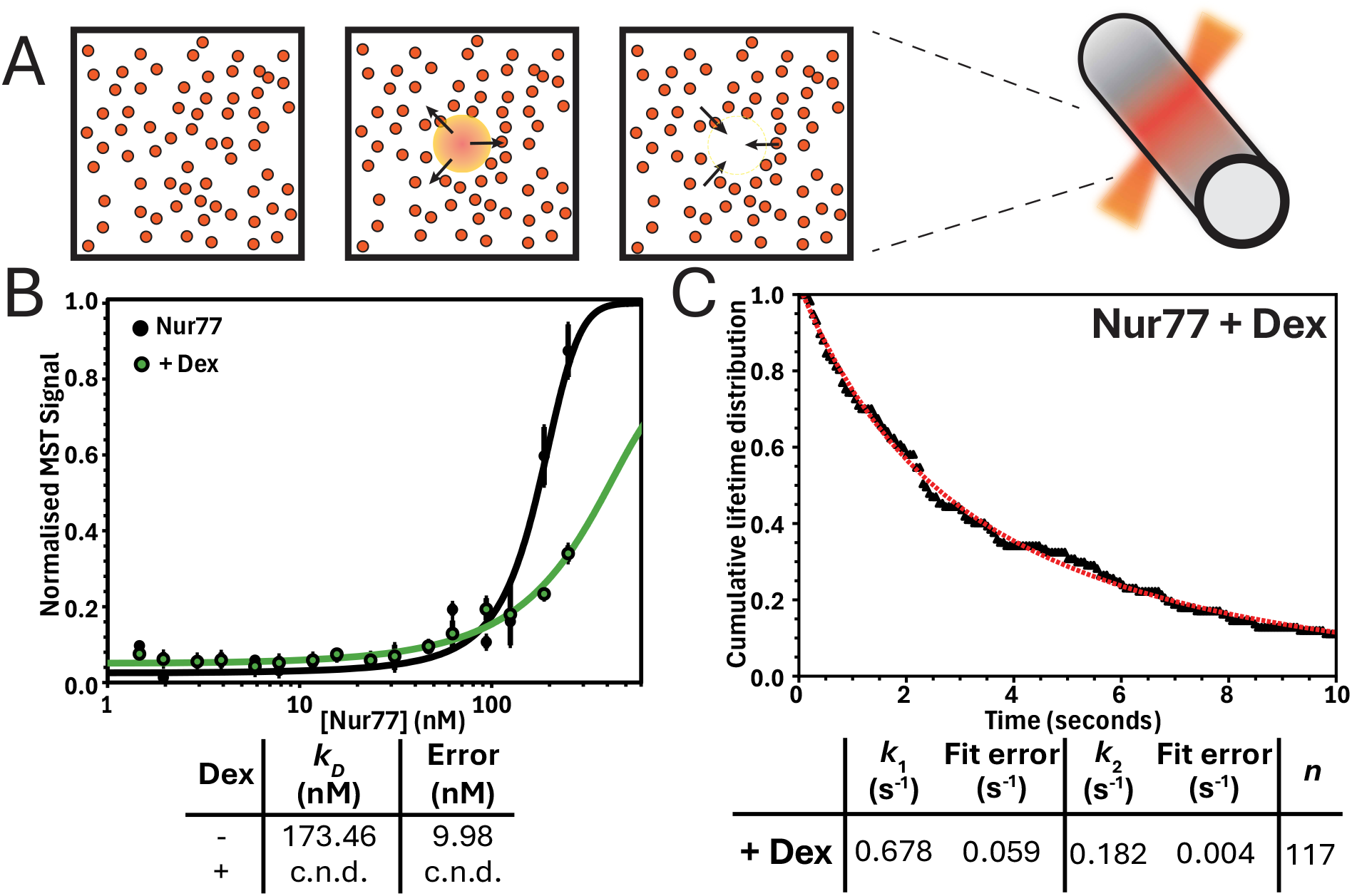
Dexamethasone (Dex) alone disrupts Nur77/DNA interactions. **(A)** Cartoon representation of an MST assay. The fluorescence emission within the capillary is continuously monitored. Upon localized heating by an infrared laser, molecules diffuse away from the heated region at a rate constant inversely correlated with their hydrodynamic size. **(B)** MST of Nur77 titrated against DNA without (black) and with (green) Dex. *k*_*D*_ and error extracted from fitting the MST values with a binding curve are represented in the table below the graph. Means ± S.E.M., *n* = 6, c.n.d. = k_D_ cannot reliably be determined. **(B)** Cumulative lifetime distribution from the single molecule tightrope assay reveals that the kinetic parameters of Nur77 with Dex is significantly altered,. Black triangles represent the data, the red dotted line the double exponential fit. Dissociation rate constants (*k*_*1*_ and *k*_2_) with fit error are shown in the table below. *n* represents the number of observations from three separate flow chambers.

Compared to traditional techniques such as electrophoretic mobility shift assays (EMSA), MST operates in a free solution without the need for immobilization or gel matrices. In our setup, the DNA is fluorescently labelled, which prevents any protein aggregates from skewing the results. Our experiments revealed that while the dissociation constant for Nur77 with unspecific DNA is174.46 ± 9.98 nmol/L, modulation by Dex reduced DNA binding, and a *k*_*D*_ could not reliably be determined (Figure 3B).

Next, we introduced Dex into the DNA tightrope assay. This revealed a similar image, where addition of Dex yielded a profoundly different outcome compared to both the Nur77 alone and the Nur77 and GR combined (Figure 3C and S4 and S5). Dex modulation suppressed the detection frequency of DNA binding, (117 events compared to 809 events of Nur77 alone) and decreased the *k*_off_ of the residual observable Nur77 (*k*_1_: 0.687 ± 0.059 s^-1^; *k*_2_: 0.182 ± 0.004 s^-1^, Figure 3B), and induced a significant population shift towards the longer-lived state (∼61% vs. ∼37% baseline). To understand this shift better, we performed subsequent MST analysis on Nur77 bound to different DNA constructs (Table S1) in the presence of Dex. This revealed that Dex strongly affected the binding of Nur77 to NurRE DNA, while increasing the *k*_*D*_ of Nur77 with NBRE DNA (Figure S6).

## Discussion

The mechanism by which the GR influences the DNA binding dynamics of Nur77 is critical in modulating transcriptional responses. Yet the molecular details of this process, and strategies to potentially modulate it, remain elusive. In this study, by combining single-molecule and biophysical analyses, we obtained several key insights that refine our understanding of this interplay. Firstly, our direct visualization of Nur77 on DNA revealed an intrinsic 3D diffusion-based search mechanism on non-specific DNA, characterized by transient interactions. This observation provides a fundamental understanding of Nur77, revealing a predominant ‘probing’ mode rather than extensive linear tracking along the DNA backbone, which may contribute to its ability to rapidly respond to cellular signals by efficiently scanning the genome.

We observed that the interactions of Nur77 are best described as consisting of two kinetic populations, a kinetic behaviour that we have previously identified for the androgen receptor (AR)^49^. *in vivo*, one of the characteristics of Nur77 is its ability to bind DNA as a monomer at the NGFI-B-responsive element (NBRE; AAGGTCA), unique for a nuclear receptor, as well as a homo- or heterodimer (with other NR4A members) to Nur-responsive elements (NurRE; TGATATTTn_6_AAATGCCA) in the promotor regions of target genes. This versatility underlies the involvement of Nur77 in fundamental processes like apoptosis, metabolism, and immune function^7,19,50^. The λ phage DNA construct we use in our tightrope assay lacks full sites of the NurRE but does contain a number of half motif sequences as well as a few NBRE-like motifs. The presence of these (half) sites may explain the observation of two populations, where Nur77 bound to a (partial) motif could increase the dwell time on DNA. Observations of longer dwell times on DNA *in vitro* are proposed to be caused by the presence of specific DNA motifs^51^. Structural studies on other member of the nuclear receptor superfamily have revealed a DNA probing mechanism where a monomer binds to half a motif, inducing a conformational change that allows the second monomer to bind^52–54^. It was furthermore revealed that in the AR this conformational change induced helicity in the N-terminal domain, which would allow co-regulators to interact with the receptor, structurally linking DNA interaction with regulating gene expression^55^. Given the immense size of the genome that these receptors must scan, and the relatively short DNA recognition motifs, it is plausible that this is a common mechanism amongst nuclear receptors, and may be a structural explanation for the population of binding events that we observed. More experiments, including structural elucidation of the Nur77 DBD in complex with DNA, are needed to confirm this model.

We found that interaction with GR significantly alters these dynamics by markedly stabilizing Nur77’s association with DNA. This was evidenced by an increased average residence time, primarily driven by a reduction in the k_off_ of the short-life population of Nur77. This is in line with previous experimental evidence, where direct protein-protein interactions lead to repression of Nur77^31^. Since experimental evidence has revealed that GR does not bind to the response elements of Nur77^27^, it is most likely that GR physically sequesters Nur77 at other sequences, like the glucocorticoid response elements (GREs; AGAACAn_3_TGTTCT). Just like the NurRE, there are no full GREs along the lambda DNA, but there are approximately 40 half sites. Such modulation of Nur77’s DNA dwell time by the GR protein itself offers a direct biophysical mechanism through which GR could broadly influence Nur77’s transcriptional output. By prolonging Nur77’s interaction with DNA, GR might enhance its capacity to engage with co-regulators or the basal transcriptional machinery at specific loci, or conversely, sequester Nur77 at non-productive sites, thereby providing a nuanced means of controlling its activity beyond simple steric hindrance at response elements. Cross referencing transcription factor binding energetics on DNA with transcriptional output revealed that dwell time is a regulatory factor, that is influenced through several mechanisms including transcription factor concentration, co-regulators, DNA sequence and genome structure^56–60^.

Our investigation furthermore uncovered a striking direct effect of Dex on Nur77’s DNA interactions. Independent of GR, Dex severely attenuated the frequency of Nur77 binding events, while dramatically increasing the stability of DNA binding and shifting the population towards the longer-lived state for those few interactions that did occur. Looking at the differences between different DNA motifs, this kinetic profile strongly suggests that Dex directly impacts Nur77 by severely hindering its DNA binding ability and/or locking the remaining NBRE bound Nur77 into a hyper-stable (monomeric) conformation. These findings are particularly intriguing, as they suggest that the widely recognized antagonism between GR and Nur77 signalling pathways may be compounded by direct, GR ligand-specific effects on Nur77’s fundamental ability to bind DNA. This introduces a previously unappreciated layer of complexity, implying that the cellular concentration and availability of glucocorticoids may directly influence Nur77’s genomic targets and residence times, potentially requiring a re-evaluation of glucocorticoid action in Nur77-regulated processes, independent of GR’s canonical functions.

In conclusion, we present the first mechanistic characterisation of Nur77-DNA interactions. The combination of real-time single molecule and biophysical analysis revealed a characteristic behaviour of Nur77, and how this behaviour is affected by another nuclear receptor and a common synthetic glucocorticoid *in vitro*. Important questions remain. Like most nuclear receptors, Nur77 is a complex multidomain protein, whose individual regions serve diverse and context-dependent functions. While structural and functional characterization of the DBD and LBD of Nur77 exists (Figure 1C), the role of the NTD remains elusive and may provide vital information on the effects of the protein-protein and/or protein-ligand interactions we have observed here. Moving forward, the distinct modulatory effects observed *in vitro* require validation within the native cellular environment. Utilizing advanced live-cell imaging techniques (fluorescence correlation spectroscopy (FCS), single molecule tracking or fluorescence recovery after photobleaching (FRAP)) can assess the nuclear mobility and chromatin residence times of Nur77 under conditions where it interacts with GR or is exposed solely to dexamethasone. Furthermore. given the increasing evidence that transcription factors operate within transcriptional biomolecular condensates^61^, it will be important to investigate the formation and dynamics of Nur77 containing condensates, and whether this is influenced by GR-Nur77 interactions or by the specific presence of dexamethasone. These experiments will determine if the biophysical mechanisms we describe here translate into functional changes in gene expression regulation within chromatin.

## Acknowledgements

We thank Thomas Schmidt (Universiteit Leiden) for providing access, technical assistance and discussions for the fluorescent microscopy experiments and the Macromolecular biochemistry group at the Leiden Institute for Chemistry for access and technical assistance to the Microscale Thermophoresis facilities.

