## Supplementary Information for "Modulation of Nur77-DNA Interactions by the Glucocorticoid Receptor"

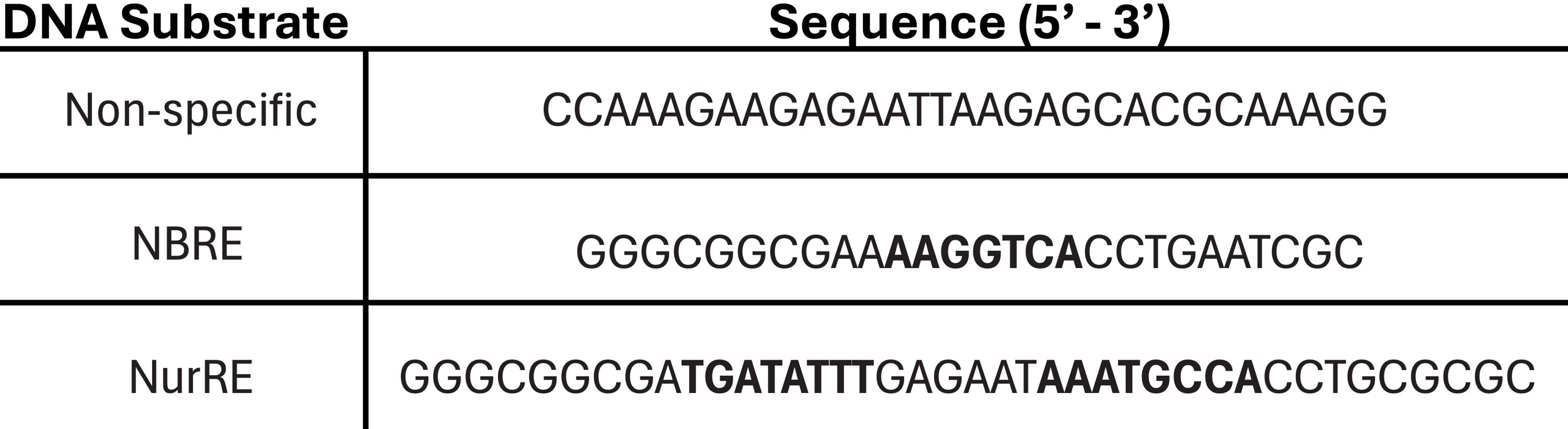
**Supplementary Tables**

**Table S1** DNA sequence of the construct used in MST.

**Supplementary Figures**

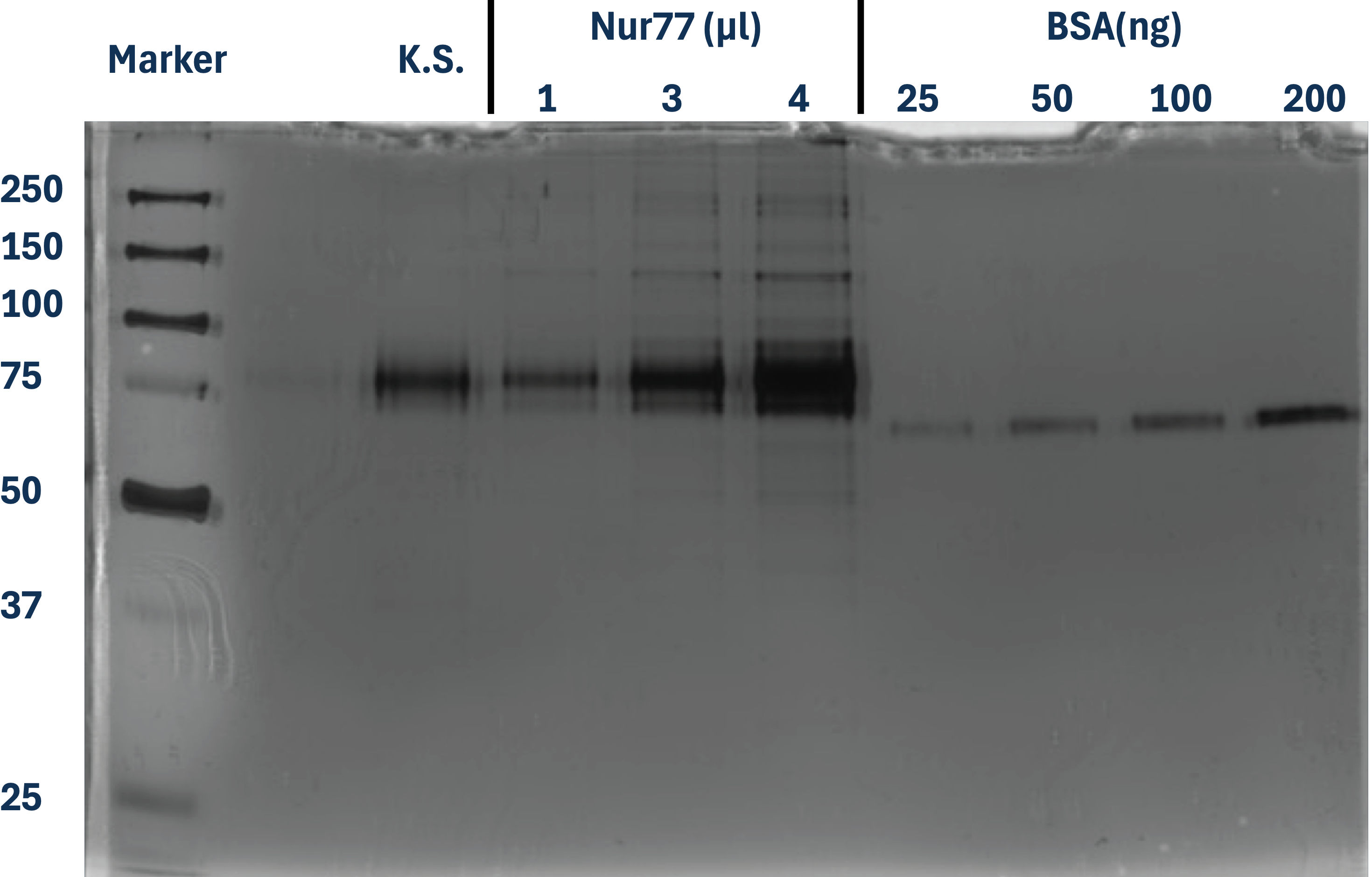

**Figure S1** A 10% SDS-PAGE Coomassie-stained gel. This gel was used to analyse the purity and concentration of Nurr77. Increasing volumes (1, 2, and 4 µl) of the Nurr77 sample were loaded alongside known sample of Nur77 (K.S.) and a quantitative standard curve of Bovine Serum Albumin (BSA) (25, 50, 100, and 200 ng). Concentration of Nur77 was determined to be 211 ng/µl. The leftmost lane contains a molecular weight ladder.

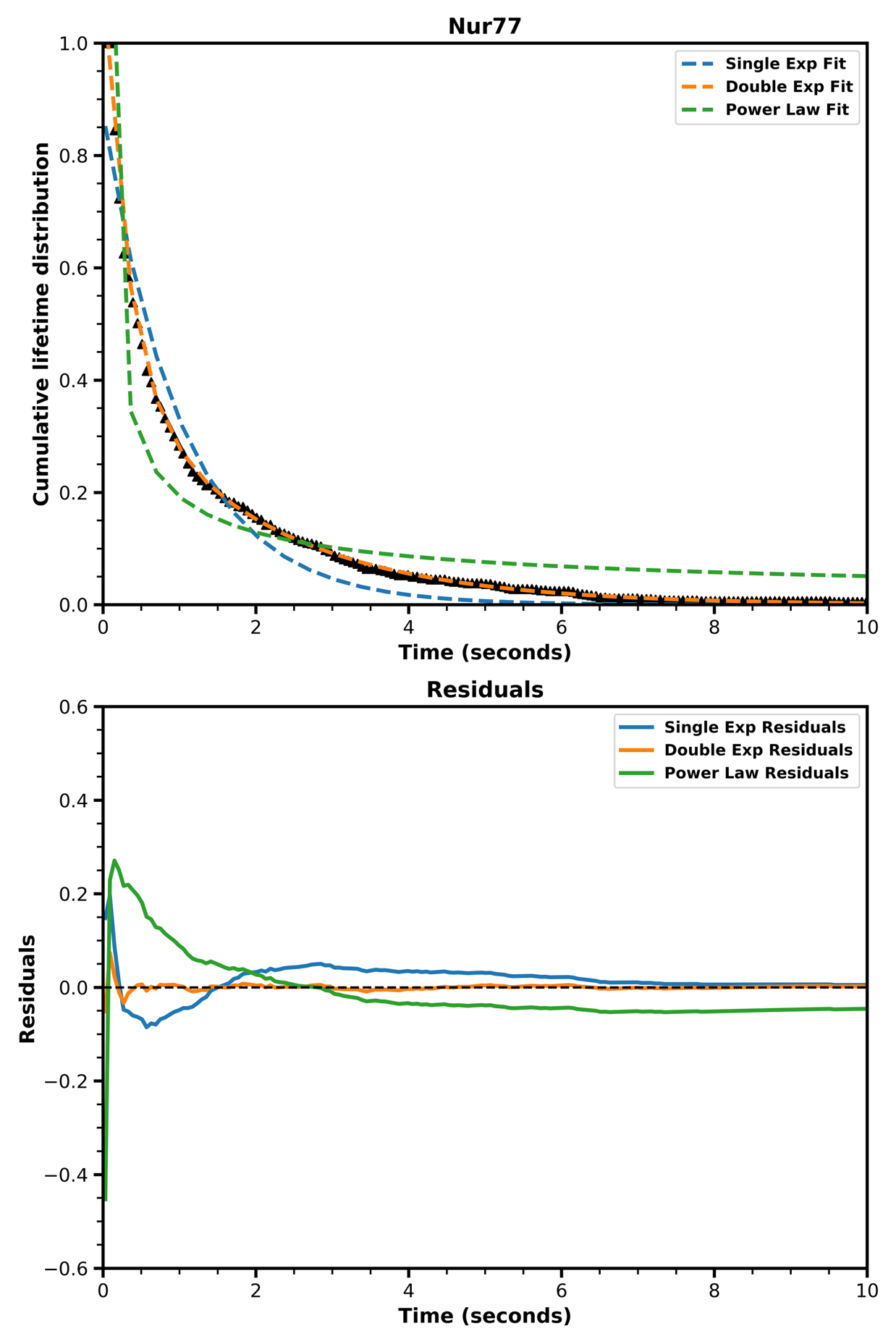
**Figure S2** Single-exponential, double-exponential and power-law fitted to the cumulative lifetime distribution of Nur77 on DNA, with corresponding residuals plot. Black triangles represent the data, blue dotted line the single exponential fit, the orange dotted line the double exponential fit, green dotted line the power law fit.

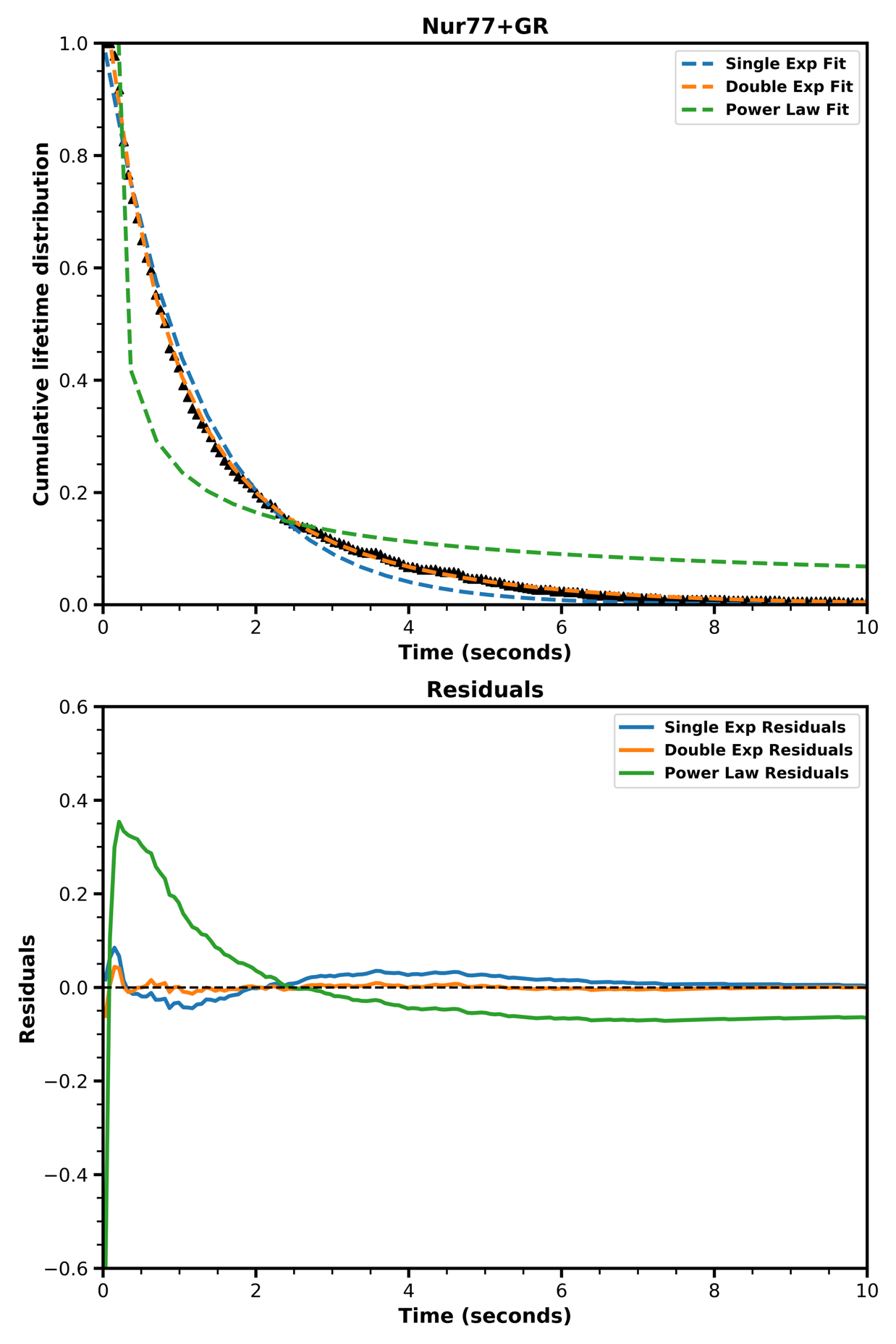
**Figure S3** Single-exponential, double-exponential and power-law fitted to the cumulative lifetime distribution of Nur77 with the glucocorticoid receptor (GR) on DNA, with corresponding residuals plot. Black triangles represent the data, blue dotted line the single exponential fit, the orange dotted line the double exponential fit, green dotted line the power law fit.

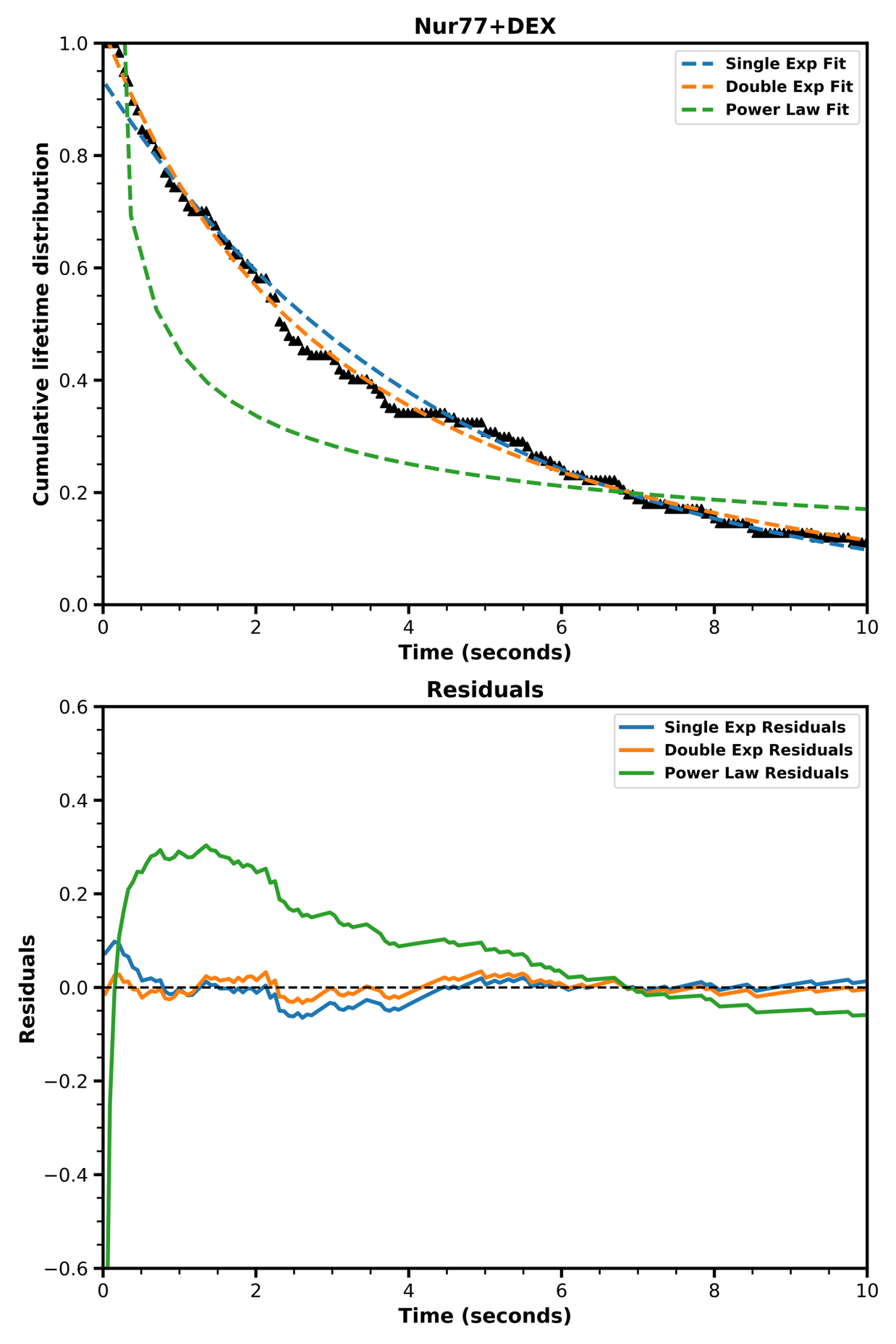
**Figure S4** Single-exponential, double-exponential and power-law fitted to the cumulative lifetime distribution of Nur77 with dexamethasone (Dex) on DNA, with corresponding residuals plot. Black triangles represent the data, blue dotted line the single exponential fit, the orange dotted line the double exponential fit, green dotted line the power law fit.

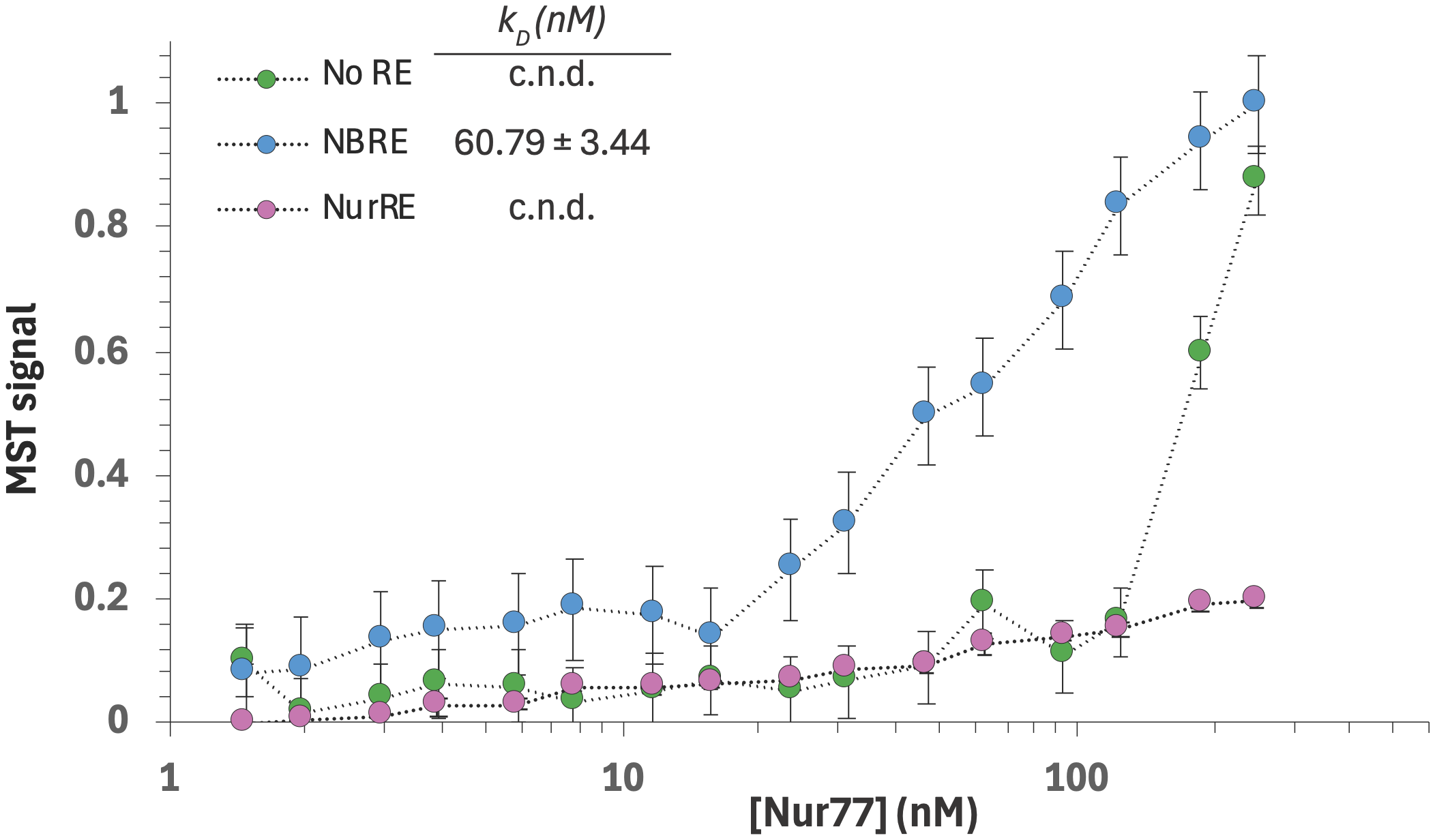

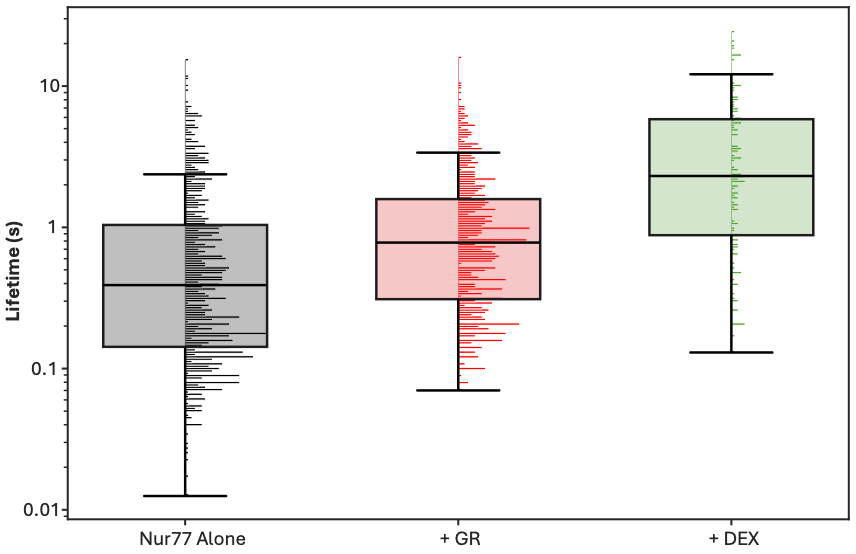
**Figure S5** Boxplot with attached normalised histograms of the molecular lifetimes (in seconds) reveal that (Dex activated) GR and Dex significantly increase Nur77 lifetimes on DNA (Mann-Whitney U test *p* < 0.001).

**Figure S6** Normalised MST results of Nur77 binding to DNA sequences without a recognition sequence (No RE), the NBRE or the NurRE sequences. Error bars indicate the standard deviation of three independent measurements. Dashed lines are lines to guide the eye. *k_D_* values and errors were extracted from the binding fits. c.n.d. = *k_D_* cannot reliably be determined.
